# α-Synuclein facilitates dopamine release during burst firing of substantia nigra neurons *in vivo*

**DOI:** 10.1101/2020.06.10.145110

**Authors:** Mahalakshmi Somayaji, Stefano Cataldi, Robert H. Edwards, Eugene V. Mosharov, David Sulzer

## Abstract

α-Synuclein is expressed at high levels at presynaptic terminals, but defining its role on neurotransmission under physiologically-relevant conditions has proven elusive. We report that α-synuclein is responsible for a rapid facilitation of dopamine release during action potential bursts *in vivo*. This occurs in tandem with a far slower stimulus-dependent depression, appears to be independent of the presence of β- and γ-synucleins or effects on presynaptic calcium and is consistent with a role for synucleins in the enhancement of synaptic vesicle fusion and turnover. The results indicate that the presynaptic effects of α-synuclein are dependent on specific patterns of neuronal activity.

## Introduction

α-Synuclein (α-Syn) is an abundant and highly conserved cytosolic neuronal protein initially identified as a constituent of cholinergic presynaptic terminals that innervate the electric organ of the torpedo electric fish (Maroteaux et al., 1988) and presynaptic inputs to cerebellar Purkinje cells (Nakajo et al., 1990). α-Syn has been reported to constitutes 0.5 – 1% of the total protein in human and rat brain (Iwai et al., 1995). Two closely related genes, β- and γ-synuclein (β-Syn: γ-Syn) were identified in brain and peripheral organs (Jakes et al., 1994). α-Syn was identified as a major constituent of disease-related protein aggregates (Ueda et al., 1993), including the Lewy body inclusions in Parkinson’s disease and other neurodegenerative disorders (Spillantini et al., 1998). Rare α-Syn mutant alleles (Polymeropoulos et al., 1997) and SNCA gene duplications and triplications (Singleton et al., 2003) cause genetic forms of Parkinson’s disease.

Synucleins are amphipathic molecules that bind to acidic phospholipids on synaptic vesicles and other highly curved membranes (Davidson et al., 1998). Following synaptic vesicle fusion, fluorescently-labelled α-Syn dissociates from the vesicles to disperse from sites of exocytosis (Fortin et al., 2005). In common with other proteins that aggregate in neurodegenerative diseases, including tau, amyloid, and TDP-43, the normal actions of the synucleins on neurotransmitter release have remained poorly understood. To date, only modest effects on neurotransmitter release have been reported in studies of synuclein-deficient animals, consisting of a small increase in the time required for presynaptic recovery following a stimulus, and an increase in dopamine release from “triple knockout” (TKO) mice lacking α-,β- and γ-Syn (Abeliovich et al., 2000; Anwar et al., 2011; Logan et al., 2017; Senior et al., 2008; Sulzer and Edwards, 2019; Yavich et al., 2004).

It was recently shown that the synucleins act to enhance the rate of fusion pore dilation during secretory vesicle exocytosis (Logan et al., 2017). This effect was deduced from the slowed exocytosis of neuropeptides from large dense core vesicles in SynTKO mice. The neuropeptides are larger than classical neurotransmitters and take longer to diffuse from the vesicle lumen during exocytosis than classical small molecule neurotransmitters during transient synaptic vesicle fusion events (~50 μs for dopamine in synaptic vesicles) (Staal et al., 2004). It remains unclear how a role for synuclein at the fusion pore might affect neurotransmitter release from synaptic vesicles.

We conjectured that if synucleins promote vesicle pore dilation, they may selectively enhance neurotransmitter release during bouts of high neuronal activity. This might occur by enhancing the clearance of vesicle membrane from presynaptic active zones, so that other synaptic vesicles might “refill” the active zone. The role of such mechanism would be particularly important at synapses that undergo bursts in the midst of ongoing tonic activity, such as from release sites on axons of ventral midbrain dopamine neurons where limited levels of presynaptic scaffolding proteins may constrain the number of active zones available for synaptic vesicle fusion (Liu et al., 2018a; Pereira et al., 2016). The acute striatal slice preparation provides a useful system for dissecting effects on dopamine release and reuptake, but lacks important attributes for differentiating the effects of axonal physiological stimulus patterns, as the dopamine cell bodies are absent and the axons do not receive physiological regulation by a variety of systems, including ongoing cortical and thalamic activity, that are important *in vivo* (Sulzer et al., 2016).

To determine whether synucleins regulate release of classical neurotransmitters from synaptic vesicles during physiologically relevant tonic and phasic activity, we stimulated the cell bodies of substantia nigra (SN) midbrain neurons and characterized evoked striatal dopamine release and dopamine axonal calcium transients *in vivo* under a state of light anesthesia. The results reveal that in wild-type (WT) mice, α-Syn exerts a rapid and profound burst firing-dependent regulation of dopamine release, consistent with a more rapid turnover of synaptic vesicle membrane and access of synaptic vesicles to presynaptic active zone fusion machinery, as well as a far slower stimulus-dependent longer-term depression. This role for α-Syn may be particularly important for synapses that undergo prolonged bouts of intermittent high activity that can undergo facilitation, including monoaminergic neurons.

## Results

### Tonic and bursting activity of ventral midbrain dopaminergic neurons under isoflurane anesthesia

During awake behavior, ventral midbrain dopamine neurons exhibit tonic firing that is interrupted by bursts of activity associated with environmental stimuli, including unexpected rewards and behaviorally salient cues, as observed in rodents (Phillips and Olds, 1969) and nonhuman primates (Mirenowicz and Schultz, 1996). In mice, the tonic pacemaking activity of dopamine neurons consists of action potentials at frequencies of 1-8 Hz that are dependent on the activities of L-type calcium channels and hyperpolarization and cyclic nucleotide gated (HCN) channels (Guzman et al., 2009), while bursting is controlled by a combination of excitatory and inhibitory synaptic inputs (Paladini and Roeper, 2014).

We recorded spontaneous firing activity of SN midbrain neurons by extracellular *in vivo* recordings from wild-type mice during isoflurane anesthesia (Subramaniam et al., 2014), from which wild-type mice recover within seconds following removal of the isoflurane. These neurons displayed firing properties typical for dopaminergic neurons, including their distinctive tonic, irregular and phasic firing patterns **(Fig 1A-F)**. We then performed juxtacellular labeling of the cells next to the recording electrodes followed by post-hoc immunolabeling for tyrosine hydroxylase, which confirmed that recorded cells were dopaminergic neurons of the SN pars compacta **(Fig 1G, H)**.

**FIGURE 1:**
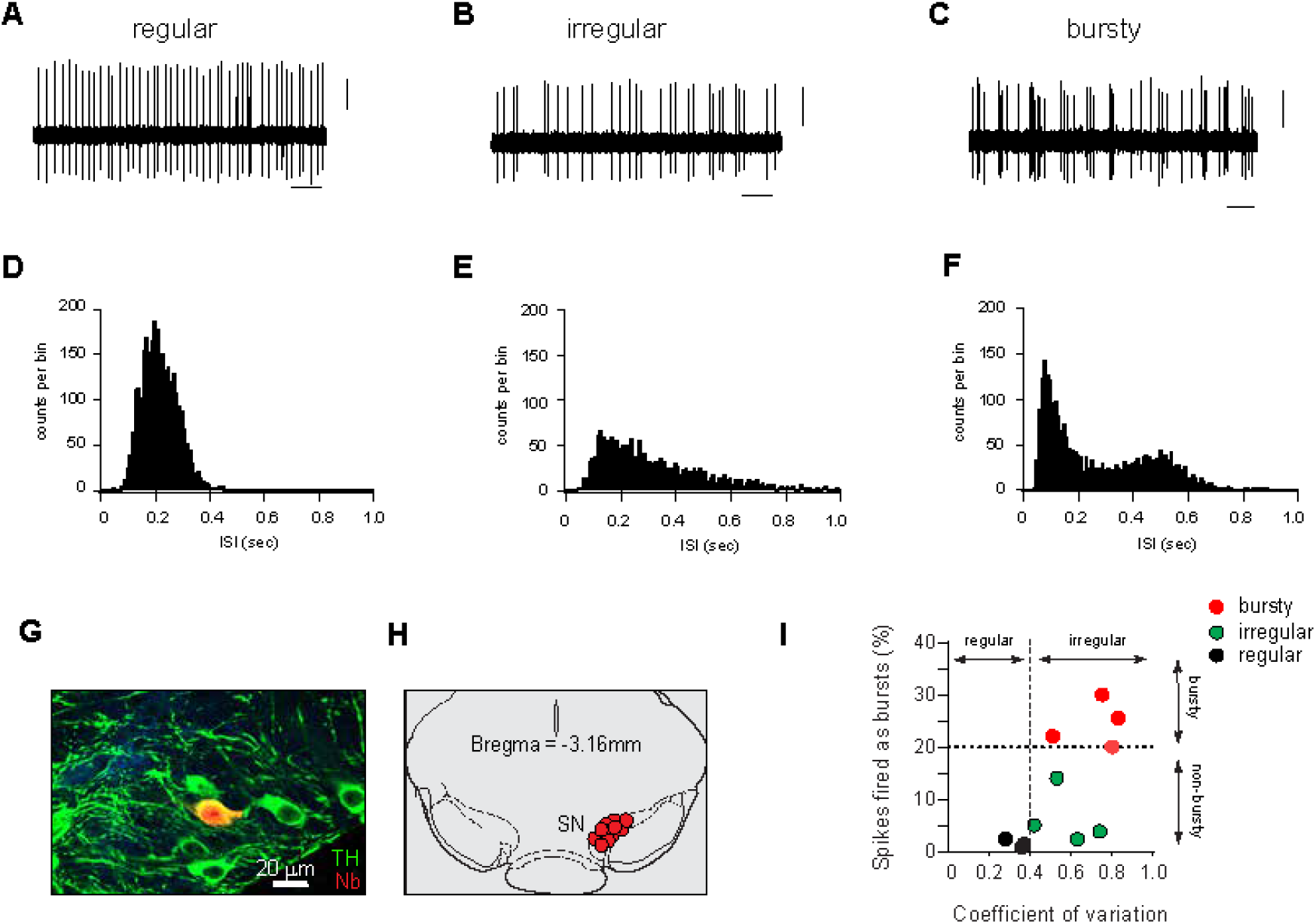
*In vivo* electrophysiological recordings from midbrain dopaminergic neurons in lightly anesthetized mice: **A-C**, Representative extracellular *in vivo* recording of spontaneous firing activity of midbrain dopamine neurons, showing examples of regular (pacemaker, A), irregular (B) and burst firing (C) patterns, respectively. Scalebar, 1 sec, 5 mV. **D-F**, Representative histograms of interspike intervals (ISI) of individual regular (D), irregular (E) and bursty (F) neurons, demonstrating the variations in the firing pattern. **G**, *In vivo* juxtacellular labeling and immunocytochemical staining showing the neurochemical identity and anatomical localization of a recorded dopaminergic neuron. Confocal laser scanning microscopic images of PFA-fixed mouse brain tissue shows neurons stained for tyrosine hydroxylase (TH, green) to label dopaminergic neurons and neurobiotin (Nb, red) to visualize a neuron near the recording micropipette. **H,** Anatomical mapping of all recorded WT DA neurons (n = 11) and their localization within the SN (coronal midbrain image adapted from the Paxinos Mouse Brain Atlas, 4^th^ ed). **I**, Scatter plot showing the coefficient of variation (mean/standard deviation) and a fraction of spikes fired as bursts (SFB) of identified midbrain dopamine neurons. Line at SFB 20% represents the threshold for neurons classified as bursty together with respective autocorrelogram-based classification. Note that ~35% of the substantia nigra dopaminergic neurons exhibit burst firing under isoflurane anesthesia.

To analyze spontaneous *in vivo* activity of these neurons, we quantified “burstiness” using a criterion defined by Grace and Bunney (Grace and Bunney, 1984), where an interspike interval (ISI) <= 80 ms defines the start of the burst and ISI >= 160 ms the end of the burst. Neurons are classified as “bursty” if the fraction of spikes fired as bursts (SFB) is greater than 20% of the total number of action potentials **(Fige 1I)**. Firing patterns of identified dopamine neurons were classified as regular or irregular based on autocorrelagrams for each neuron (see Methods). Based on the analysis, irregular patterns could also be detected as having a CV >= 0.4 (Fig 1I).

In mice undergoing light isoflurane anesthesia, the majority of dorsal striatum-projecting dopaminergic SN neurons fired tonically, while about 35% also displayed significant bursts interspersed during the tonic activity (18 % regular; 46% irregular). While we cannot rule out the possibility that light isoflurane anesthesia may completely silence some dopaminergic neurons, cells that alternated between tonic activity and silence were not observed. These firing patterns contrast with an absence of tonic activity reported in a fraction of neurons during deeper anesthesia with chloral hydrate (Koulchitsky et al., 2012).

### Characterization of stimulus parameters to elicit DA release

Since the introduction of the carbon fiber microelectrode (Gonon et al., 1980), amperometry and cyclic voltammetry have been used to measure evoked dopamine release and reuptake in anesthetized rodents (Sulzer et al., 2016). We used fast scan cyclic voltammetry (FSCV) to measure extracellular dopamine at sub-second temporal resolution.

Pioneering studies by Francois Gonon demonstrated that in the absence of reuptake blockers, stimuli that emulate the tonic activity of ventral midbrain dopamine neurons do not produce dopamine release *in vivo* that can be measured by electrochemistry (Chergui et al., 1994), which typically has a limit of detection of ~50 nM dopamine in the extracellular space (Mosharov et al., 2003; Sulzer et al., 2016). In contrast, burst firing of dopamine neurons at >10 Hz can drive dopamine levels to 1 μM or higher (Garris et al., 1997; Gonon, 1988), as a buildup of extracellular dopamine saturates the dopamine uptake transporter (DAT) (Schmitz et al., 2003; Sulzer et al., 2016).

We first confirmed that in animals under isoflurane anesthesia, no changes in extracellular striatal DA were observed in the absence of stimuli (not shown), even though the cell bodies of SN neurons innervating this area were active (Fig 1). We then stimulated SN cell bodies electrically while measuring the evoked DA release in the dorsal striatum using the placement of the stimulus electrode as for the extracellular recordings in Fig 1. To confirm the placement of the recording carbon fiber microelectrode, we coated it with a fluorescent lipophilic membrane dye, 1,1’-dioctadecyl-3,3,3’3’-tetramethyl-indocarbocyanine perchlorate (DiI). The bipolar stimulation electrode tracks in the ventral midbrain and DiI-stained recording electrode track in the dorsal striatum were clearly observed in post-fixed brain slices **(Fig 2A)**. The oxidation profile and the time course of dopamine release in the dorsal striatum is shown as a color plot on **Fig 2B**. A characteristic background-subtracted voltammogram at the maximum of the oxidation peak is consistent with the release of dopamine **(Fig 2B inset)**.

**FIGURE 2:**
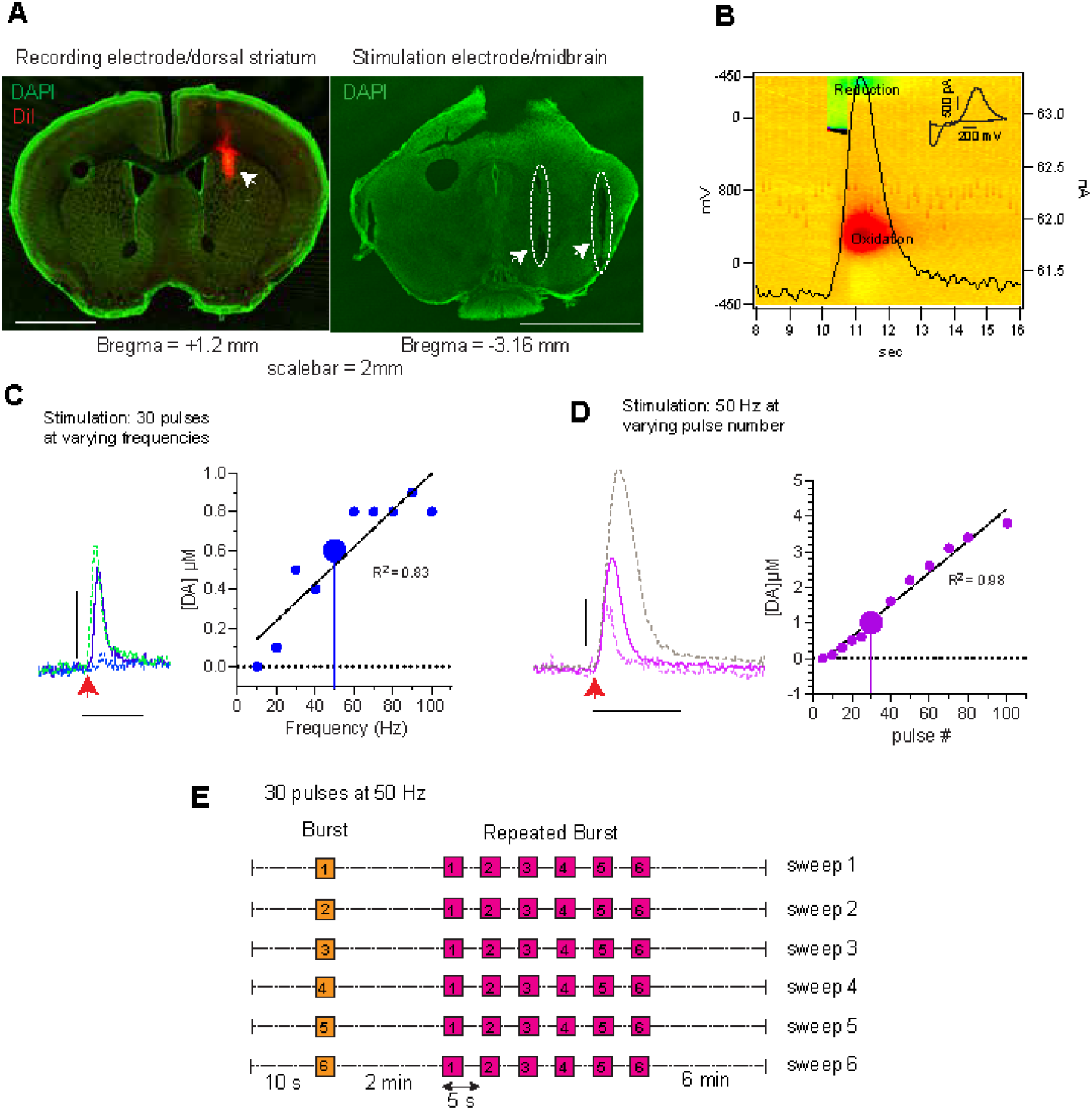
Characterization of evoked dopamine release protocol for *in vivo* anesthetized mice recordings by fast scan cyclic voltammetry. **A**, Confocal laser scanning microscope images (5x objective) of PFA-fixed coronal mouse brain sections indicating the location of recording electrode in the dorsal striatum (left, green = DAPI, red = DiI staining, arrowhead) and the bipolar stimulating electrode in the ventral midbrain (right, dashed outline and arrowheads). Scale bar = 2 mm. **B**, 3-D pseudo-color plot showing oxidation (red) and reduction (green) of DA. The time course of oxidation at 300 mV is superimposed as a black trace. Insert: voltammogram at the maximum of oxidation (11.2 sec). Scale bar = 200 mV and 500 pA. **C**, *Left*: Representative peaks of evoked dopamine release obtained by stimulating the dopaminergic cell bodies in the midbrain with a constant pulse number (30 pulses) at 20 Hz (dotted), 50 Hz (line) and 90 Hz (green, dotted) frequencies. Red arrow indicates the start of electrical train of stimuli. *Right*: Correlation between evoked dopamine release and stimulus frequency (blue, fit of the data with a linear regression *y = 0.01x + 0.05*) **D**, *Left*: Examples of evoked dopamine release following stimulation of midbrain dopaminergic neurons by 20 (dotted), 30 (line) and 60 (grey, dotted) pulses at a constant 50 Hz frequency. Red arrow indicates the start of the train of stimuli. Scalebar, Y-axis = 500 nM DA, X-axis = 5 sec. *Right*: Correlation between evoked dopamine release and pulse number (magenta, linear regression *y = 0.04x − 0.3*). The larger circles represent parameters that provide a preferred dynamic range (30 pulses at 50 Hz) used in this study. **E**, Stimulus protocol developed based on the results from C and D. Each sweep is comprised of a single stimulus train (orange, 30 pulses at 50 Hz) with a recovery period of 2 min, followed by a repeating burst stimulation (magenta), consisting of 6 stimulus trains (30 pulses at 50 Hz) every 5 sec, with a recovery period of 6 min before the next sweep. The entire protocol consists of six consecutive sweeps.

To study the dependence of striatal dopamine release on SN stimulus patterns, we compared the extracellular dopamine levels as a function of stimulus frequency at a constant pulse number (30 pulses) **(Fig 2C)**, or as a function of number of pulses at a constant stimulus frequency (50 Hz) **(Fig 2D)**.

At 10 Hz stimulation frequency, we did not resolve evoked DA release, consistent with the *in vivo* studies mentioned above in which multiple stimuli at 15 Hz were required to saturate DA uptake and measure evoked DA release (Chergui et al., 1994). Increasing the frequency from 20 to 60 Hz produced a steady increase in evoked DA release, but further increase in stimulus frequency to up to 100 Hz did not change the amplitude of DA peak.

In contrast to the saturation of DA release at higher stimulation frequencies, we found that extracellular DA evoked at 50 Hz was linear between 5 and 80 pulses. This indicates that within bursts, the same amount of DA was released per pulse, and that dopamine release recovered to a stable level within 20 msec. Based on these results, for the following experiments we selected 30 pulses at 50 Hz as a stimulus that produced a response in a linear range optimal for detecting neuromodulation.

From these characterizations, we designed a protocol for the characterization of activity-dependent dopamine plasticity (**Fig 2E**), consisting of a “single burst” (orange box, a train of 30 pulses at 50 Hz) followed by a recovery period of 2 min and a train of “repeated bursts” (1-6, magenta boxes) stimuli. The latter was comprised of six “single bursts” repeated at 5 sec intervals. A “single burst” followed by a “repeated burst” constitutes a “sweep” that was repeated six times with inter-sweep intervals of 6 minutes. This protocol provides information on both short-term and long-term effects on dopamine plasticity *in vivo*.

### Synuclein-dependent deficits in activity-dependent presynaptic recovery

We compared responses to the single burst stimuli in three mouse lines, wild-type (WT), “triple knockout” deficient for α, β and γ synucleins (SynTKO) and a line in which only α-Syn is deleted (α-SynKO) (Anwar et al., 2011). Analysis of dopamine release evoked by single bursts (**Fig 3A**, shaded area) over the six sweeps showed a sweep number-dependent decrease in WT mice to ~ 50% of initial levels (**Fig 3B, C**; one-way ANOVA within WT genotype: F_5, 36_ = 5.2, *p* = 0.001, Tukey’s multiple comparison – sweep1 vs sweep 6, *p* = 0.006). In contrast, SynTKO mice showed no significant decrease in response (one-way ANOVA within genotypes: SynTKO, F_5, 36_ = 0.073, *p* = 0.99, Tukey’s multiple comparison – not significant; α-SynKO, F_5, 24_ = 0.11, *p* = 0.99, Tukey multiple comparison – not significant. Two-way ANOVA between 3 genotypes: F_2, 96_ = 12.4, *P* < 0.0001). The sweep-number dependent decrease in dopamine release in the WT was unchanged when only single bursts were applied and trans of repeated bursts were omitted from the sweeps (**Fig 3E)**.

**FIGURE 3:**
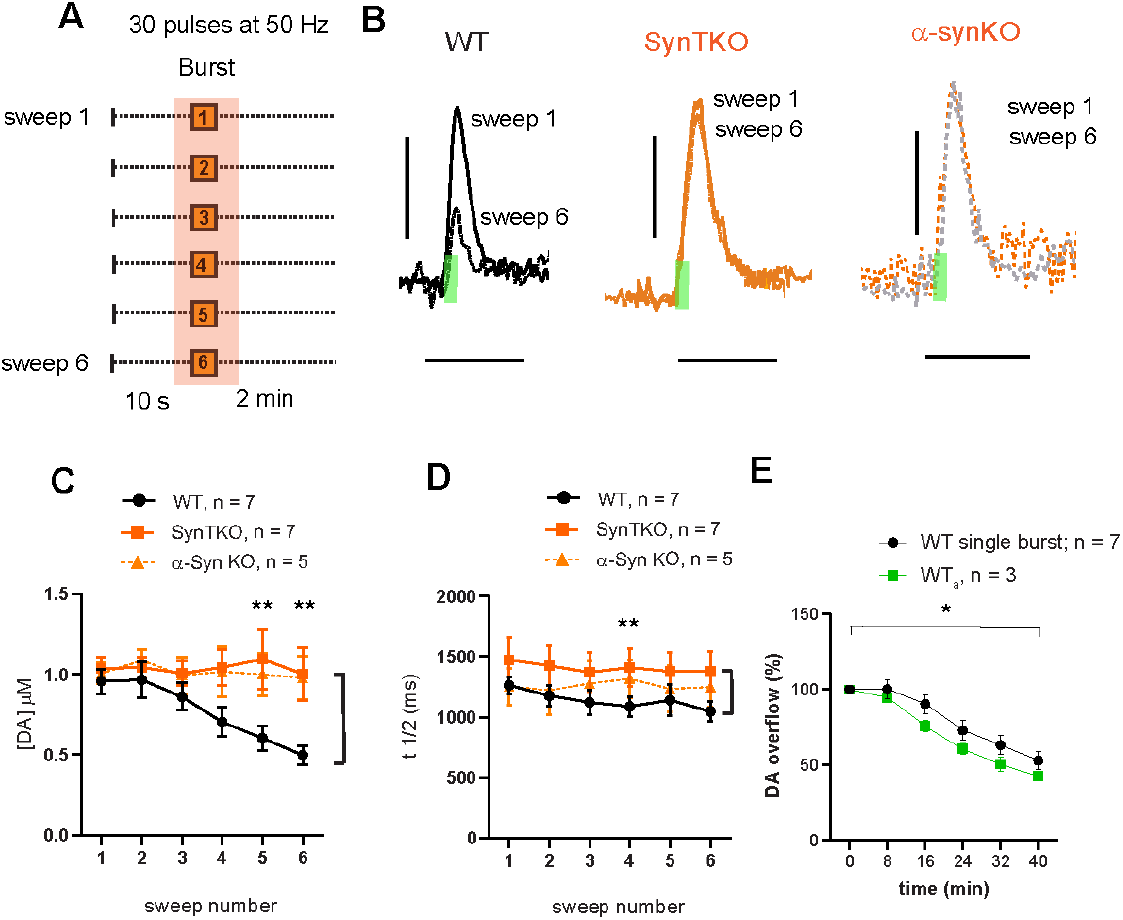
Synuclein-dependent decrease in evoked DA released during stimulation of midbrain DA neurons with single bursts with long (6 min) rest intervals. **A**, Schematic of the stimulus region used for data analysis. **B**, Evoked DA peaks following single burst stimulus (30 pulses at 50 Hz) from sweeps 1 and 6 showing the differences in DA release between wildtype (WT), synuclein triple knockout (SynTKO) and α–synuclein (α-SynKO) mice. Green bars indicate electrical stimulation duration (0.6 s). Scalebar, Y-axis = 500 nM DA, X-axis = 5 sec. **C**, DA release decreases across sweeps in WT (black) but not in SynTKO (orange) and α-SynKO (orange-broken line) mice (two-way ANOVA: Genotype: F_2, 96_ = 12.4, *P* < 0.0001, Interaction: F_10, 96_ = 1.1, *P* = 0.4. Tukey’s multiple comparison test showed significance between WT and SynTKO in burst numbers 5-6 (*P* = 0.003, 0.002 respectively), as well as WT and α-SynKO in burst numbers 5-6 (0.04, 0.009 respectively); *n*: WT = 7, SynTKO= 7, α-SynKO = 5. **D**, The t½ does not change across sweeps in any of the genotypes. There was a shorter t1/2 in WT than in SynTKO, but not α-SynKO (two-way ANOVA: WT vs SynTKO: Genotype: F_1, 72_ = 11.6, *P* = 0.001, WT vs a-SynKO: F_1, 60_ = 2.3, *P* = 0.12, n: WT = 7, SynTKO= 7, α-SynKO = 5). **E**, DA release decreased by approximately the same degree in WT mice stimulated either with single burst within a sweep (black, same data as C)) or single bursts only at 2 sec intervals (green) (two-way ANOVA: Genotype: F_1, 48_ = 5.8, *P* = 0.02, Interaction: F_5, 48_ = 0.34, *P* = 0.9. Bonferroni’s multiple comparison test did not show significance, n: WT = 7, WT_a_ = 3).

To investigate a possible role of synucleins in the regulation of DAT activity (Sidhu et al., 2004), we examined the t1/2 of the dopamine peaks. There was no change in t1/2 values within the sweeps in any of the genotypes, suggesting that dopamine reuptake was not altered during the course of the experiment (Schmitz et al., 2003; Sulzer et al., 2016). We noted a slightly longer t1/2 in the SynTKO than in WT (**Fig 3D**, see Fig Legend for statistical details).

These results indicate that α-Syn expression can cause activity-dependent depression of dopamine signaling *in vivo.* This is consistent with previous reports both *in vivo* (Yavich et al., 2004) and *in vitro* (Abeliovich et al., 2000; Anwar et al., 2011) that synuclein expression slows the time required for the recovery of evoked dopamine release.

### A role for synuclein in modulating short-term presynaptic plasticity

We then compared dopamine release kinetics during repeated bursts of stimuli (**Fig 4A**, shaded area). Similar to dopamine released in the repeated single-burst stimuli in Fig 3, the overall amount of dopamine released during each sweep, measured as the area under the curve (AUC; **Fig 4B**, shaded area), decreased with prolonged stimulation *in vivo* (one-way ANOVA within WT genotype: F_5, 36_ = 12.7, *p* < 0.0001, Tukey’s multiple comparison – sweep1 vs sweep 6, *p* < 0.0001). In contrast, SynTKO mice showed no significant decrease in response (one-way ANOVA within genotypes: SynTKO, F_5, 42_ = 0.8, *p* = 0.6, Tukey’s multiple comparison – not significant; α-SynKO, F_5, 24_ = 0.8, *p* = 0.6, Tukey multiple comparison – not significant). Consistent with the results from the single burst data (**Fig 3C**), there was a stronger depression in the WT than in the α-SynKO and SynTKO (**Fig 4E**, two-way ANOVA between genotypes: *P* = 0.04, WT, n = 7, SynTKO, n = 7 and α-SynKO, n = 5. Tukey’s multiple comparison showed significance between WT and SynTKO in sweep 6 (*P* = 0.04). See figure legend for additional statistics). Also consistent with the response to single bursts (Fig 3C), WT mice exhibited a progressive decrease in dopamine release elicited from the first burst within the repeated burst, whereas release from SynTKO and α-SynKO mice was stable (**Fig 4F**, two-way ANOVA between genotypes: *P* < 0.0001, WT, n = 7, SynTKO, n = 7 and α-SynKO, n = 5. Tukey’s multiple comparison showed significance between WT, SynTKO and α-SynKO in sweeps 5 and 6).

**FIGURE 4:**
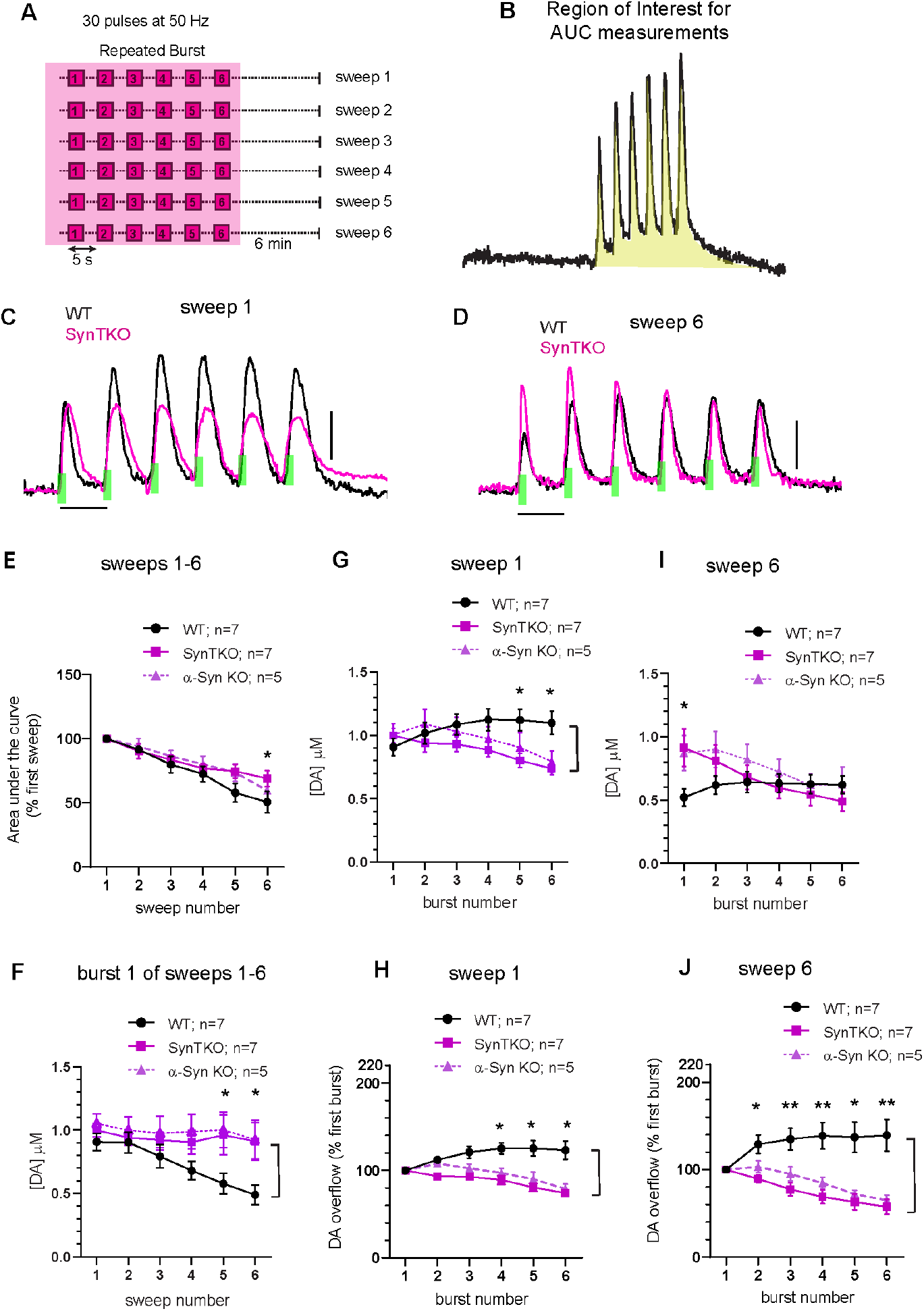
Synuclein-dependent facilitation of DA release during short-interval (5 sec) burst stimuli. **A-B,** Schematic of the regions used for data analysis (shaded). **C-D,** Representative recordings of evoked DA release resulting from repeated burst stimulations during sweep 1 (**C**) and sweep 6 (**D**). WT (black) mice demonstrate short term intra-burst potentiation, while SynTKO (magenta) mice show short-term depression in vivo. Green bars indicate the duration (0.6 s) of electrical stimulation., Y-axis = 500 nM DA, X-axis = 5 sec.**E,** Depression of overall DA release during the entire burst duration measured as total AUC (as shown on B) across sweeps 1 to 6. (Two-way ANOVA: Genotype: F_2, 96_ = 3.4, P = 0.04, Interaction: F_10, 96_ = 0.8, P = 0.7. Tukey’s multiple comparison test showed significance in sweep 6 between WT and SynTKO, P =0.04). **F,** DA release decreases across sweeps in WT but not in SynTKO and α-SynKO mice in the first burst of repeated burst protocol (two-way ANOVA: Genotype: F_2, 96_ = 11.6, P < 0.0001, Interaction: F_10, 96_ = 0.79, P = 0.6. Tukey’s multiple comparison test showed significance between WT and SynTKO in burst numbers 5-6 (P = 0.02, 0.007 respectively) and WT and α-SynKO between burst numbers 5-6 (P = 0.02, 0.01 respectively). Note the similarity to single burst changes on Fig 3C. **G and I,** Facilitation of dopamine release in WT and depression of dopamine release in the SynKO animals during the repeated burst (**Sweep 1, G:** Two-way ANOVA: Genotype: F_2, 96_ = 7.9, P = 0.0006, Interaction: F_10, 96_ = 1.5, P = 0.15. Tukey’s multiple comparison test showed significance between WT and SynTKO in burst numbers 5-6 (P = 0.01, 0.004 respectively) and WT and α-SynKO between burst number 6 (P = 0.03); **Sweep 6, I:** Two-way ANOVA: Genotype: F_2, 96_ = 3.6, P = 0.03, Interaction: F_10, 96_ = 1.3, P = 0.2. Tukey’s multiple comparison test showed significance between WT and SynTKO in burst number 1 (P = 0.006) and WT and α-SynKO between burst number 1 (P = 0.03). **H and J**, Same data as **G** and **I,** normalized to the first peak of each sweep. SynTKO and α-SynKO mice show intra-sweep depression in DA release, compared to WT, which shows DA release facilitation across bursts (**Sweep 1, I:** Two-way ANOVA: Genotype: F_2, 96_ = 49.8, P < 0.0001, Interaction: F_10, 96_ = 3.7, P = 0.0003. Tukey’s multiple comparison test showed significance between WT and SynTKO in burst numbers 2-6 (P = 0.04, 0.0008, <0.0001, <0.0001, <0.0001 respectively) and WT and α-SynKO between burst numbers 4-6 (P = 0.003, 0.0001, <0.0001 respectively). **Sweep 6, J:** Two-way ANOVA: Genotype: F_2, 96_ = 49.8, P < 0.0001, Interaction: F_10, 96_ = 2.8, P = 0.004. Tukey’s multiple comparison test showed significance between WT and SynTKO in burst numbers 3-6 (P = 0.01, 0.0002, <0.0001, <0.0001, <0.0001 respectively) and WT and α-SynKO between burst numbers 4-6 (P = 0.03, 0.002, 0.0001, <0.0001 respectively)). Data points represent mean +/− SEM; n= 7 for WT (black), 7 for SynTKO (magenta) and 5 for α-SynKO (magenta − broken line).

Remarkably, within each train of repeated bursts, the WT mice displayed a profound facilitation of evoked dopamine release, while both SynTKO and α-SynKO mice showed a depression of release (**Fig 4C, G** and **H**; two-way ANOVA between genotypes: **4G**, *P* = 0.0006, **4H**, *P* < 0.0001, WT, n = 7; SynTKO, n= 7; α-SynKO, n = 5). The difference between the relative facilitation of dopamine release in the WT and depression in the SynTKO and a-Syn KO increased even further in the later sweeps (**Fig 4D, I and J**; two-way ANOVA between genotypes: **4I**, *P* = 0.03, **4J**, *P* < 0.0001, WT, n = 7, SynTKO, n = 7, α-SynKO, n = 5).

To our knowledge, these data are the first to demonstrate a role for synucleins in facilitating neurotransmitter release, and that this occurs during bouts of high presynaptic activity.

### Synuclein-dependent dopamine plasticity *in vivo* is not due to residual presynaptic calcium

The classical model of presynaptic facilitation it that it is due to an increased accumulation of residual calcium during closely-spaced repetitive stimuli (Katz and Miledi, 1968). We examined whether this mechanism might explain α-Syn-dependent presynaptic facilitation. We compared evoked calcium transients from dopamine axons in WT and α-SynKO mice by recording GCaMP6f signals from dopamine axons by fiber photometry in the dorsal striatum using the same coordinates and stimulus protocols detailed above for electrophysiology and FSCV. As SynTKO and α-SynKO mice displayed similar evoked DA release properties, we used the α-SynKO line for these experiments. The area under the curve (AUC) of the GCaMP signal depended linearly on the number of applied stimuli (**Supplemental Fig 1**), and signals returned to baseline in < 2 sec after bursts (**Fig 5** and **Supplemental Fig 1**).

**FIGURE 5:**
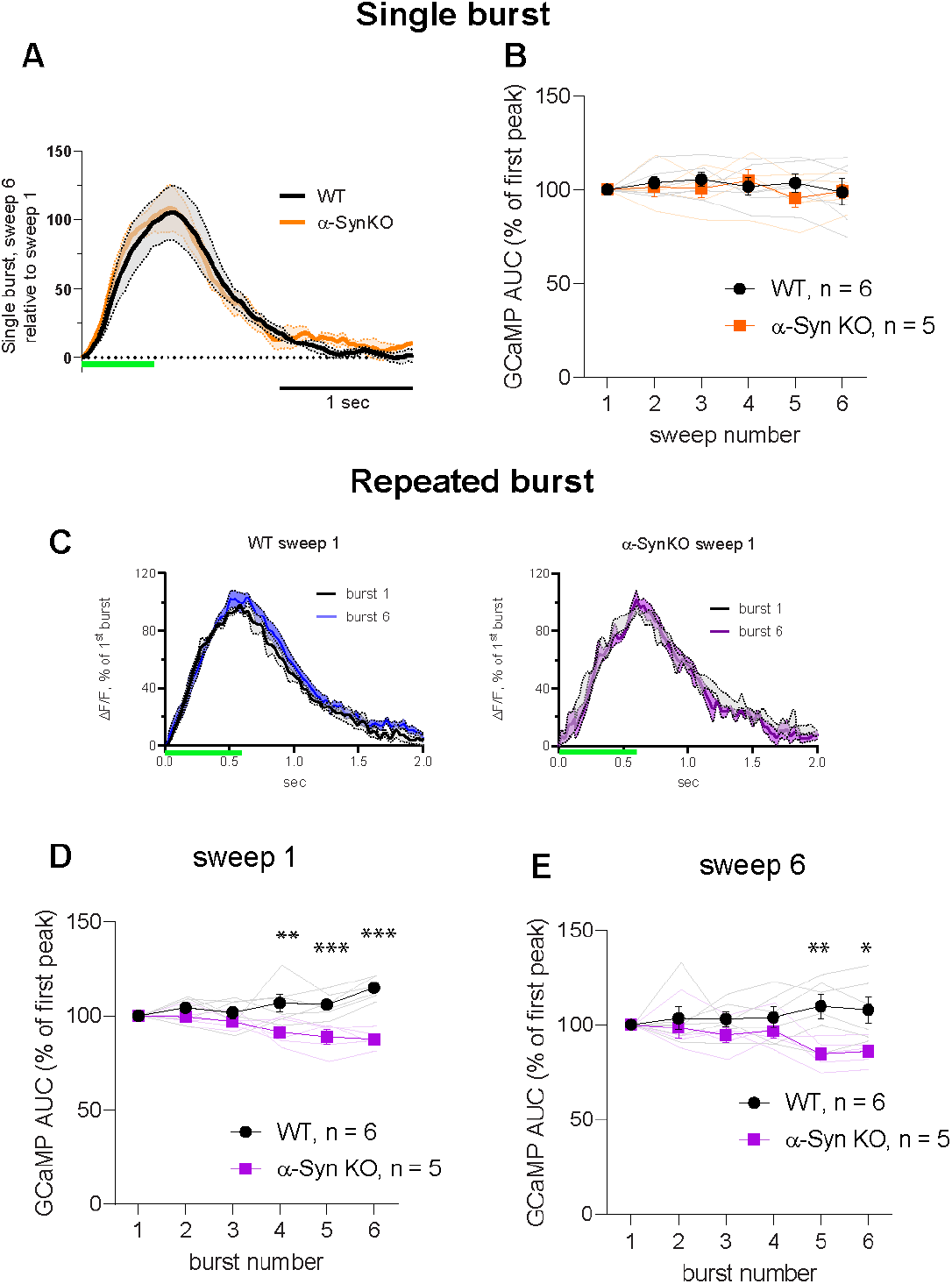
Calcium signals in response to burst stimuli *in-vivo*. **A**, Changes in evoked GCaMP6f signal transients during single burst stimulation in WT (black) and ⍰-SynKO (orange) mice represented as averaged sweep 6 fluorescence relative to sweep 1 signal. No change in the amplitude or t1/2 was detected. **B**, No change in the mean AUC of GCaMP responses at any sweep number was detected between the WT (black) and ⍰-SynKO (orange) mice (two-way ANOVA: Genotype: F_1, 54_ = 0.5, *P* = 0.5, Interaction: F_5, 54_ = 0.4, *P* = 0.9. Bonferroni’s multiple comparison test did not show significance, *n*: WT = 6, α-SynKO = 5). Faint traces show recordings from individual mice, while traces in bold represent mean+/−SEM. **C**: Changes in GCaMP6f transients in sweep 1 represented as averaged signals at burst 6 normalized to burst 1 in WT and ⍰-SynKO mice. While the amplitude of evoked Ca^2+^ signal remained the same between WT (black) and ⍰-SynKO (magenta) mice, there was a small increase in the t_1/2_ of the 6^th^ GCaMP6f transient in WT neurons. Green bars indicate stimulus duration.**D and E**: AUC of GCaMP transients in sweep 1 (D) in sweep 6 (F) was significantly higher in WT than ⍰-SynKO animals. *Sweep1*, two-way ANOVA: Genotype: F_1, 54_ = 51.5, *P* < 0.0001, Interaction: F_5, 54_ = 6.7, *P* < 0.0001. Tukey’s multiple comparison test showed significance between WT and α-SynKO in burst numbers 4 - 6 (*P* = 0.001, 0.0004, <0.0001 respectively). *Sweep 6*, two-way ANOVA: Genotype: F_1, 54_ = 16.7, *P* = 0.0001, Interaction: F_5, 54_ = 2.1, *P* = 0.08. Tukey’s multiple comparison test showed significance in bursts 5-6 (*P* = 0.003, 0.01)). *n*: WT = 6, α-SynKO = 5. The error bars in the plots represent SEM.

The averaged calcium transients evoked by single bursts of stimuli identical to those in Fig 3A are shown in **Fig 5A**. No difference in GCaMP signal response was observed between WT and α-SynKO mice (**Fig 5B**, repeated measures analysis using the mixed effects model, genotype: *P* = 0.7, Bonferroni’s multiple comparisons test did not show significance).

The responses of GCaMP6f signals to train burst stimuli as in Fig 4A are shown in **Fig 5C-E**. We observed a larger signal in the later bursts within sweeps in WT than α-SynKO animals, as seen from a small increase in t1/2 of the GCaMP peak. There was however no significant increase in signal in WT, particularly by sweep 6, where facilitation of DA within bursts was observed, and no buildup due to residual calcium. While the interpretation of the GCaMP6f signals has technical limitations (see Discussion and Supplemental Figure 1), differences in the accumulation of residual calcium appears unlikely to underlie the differences in dopamine release between WT and α-SynKO lines.

## Discussion

We examined the effects of synuclein on evoked dopamine release, reuptake, and axonal calcium transients in the dorsal striatum of mice lightly anesthetized with isoflurane. Our experimental conditions were designed to emulate the normal physiological patterns of ventral midbrain dopamine neurons activity during awake behavior, in which bursts are superimposed over pace-making activity. The comparison between WT and synuclein deficient lines revealed a profound activity-dependent role for α-Syn in the facilitation of neurotransmitter release *in vivo*. As α-Syn expression results in a facilitation of dopamine release within bursts while α-SynKO and TKO exhibit a depression, α-SynKO expression can reach a two-fold relative facilitation. This presynaptic facilitation is particularly salient for neurotransmitter systems such as monoamines that engage in extrasynaptic overflow and rely on intermittent activity bursts (Sulzer et al., 2016).

As the effects of α-SynKO and SynTKO on dopamine release *in vivo* were similar, α-Syn is apparently responsible for both the rapid facilitation and a slow depression of evoked dopamine release, and that β-Syn and γ-Syn do not compensate for the loss of α-Syn at these synapses. This does not rule out a role for other synucleins in other synapses or for these synapses under other stimulus protocols.

α-Syn-dependent facilitation of dopamine is reminiscent of the classical presynaptic paired pulse facilitation extensively characterized at the neuromuscular junction and glutamatergic pyramidal synapses. These forms of facilitation, which are characterized at far shorter stimulus intervals, are thought to originate from low initial release probability (Katz and Miledi, 1968) and a buildup of residual presynaptic calcium that enables the fusion of additional synaptic vesicles. We observed a slightly larger increase in GCaMP6f signal during trains of repeated bursts in WT than α-SynKO, but no evidence for a buildup of residual calcium between successive sweeps. It thus appears unlikely that α-Syn-dependent facilitation results from the classical mechanism of presynaptic facilitation. A note of caution is warranted, however, as the recording techniques do not indicate calcium changes within axons or may lack the sensitivity or temporal resolution to identify differences between the genotypes. Further experimentation and technology development will be needed to determine more subtle or local synuclein-dependent changes in dopaminergic axon calcium responses, if any.

An alternate mechanism for α-Syn-dependent presynaptic facilitation is suggested by a role for synucleins in facilitating secretory vesicle exocytosis by enhancing the rate of dilation and collapse of the vesicle membrane during fusion at active zones (Logan et al., 2017; Sulzer and Edwards, 2019). Multiple studies indicate that activity-dependent increase of presynaptic calcium promotes refilling of competent synaptic vesicle/active zone complexes (Byczkowicz et al., 2018; Malagon et al., 2020). A synuclein-dependent enhancement of this refilling rate initiated during burst firing-induced calcium entry could enable facilitation by a more rapid replacement of synaptic vesicles/active zone complexes (**Fig 6**).

**FIGURE 6:**
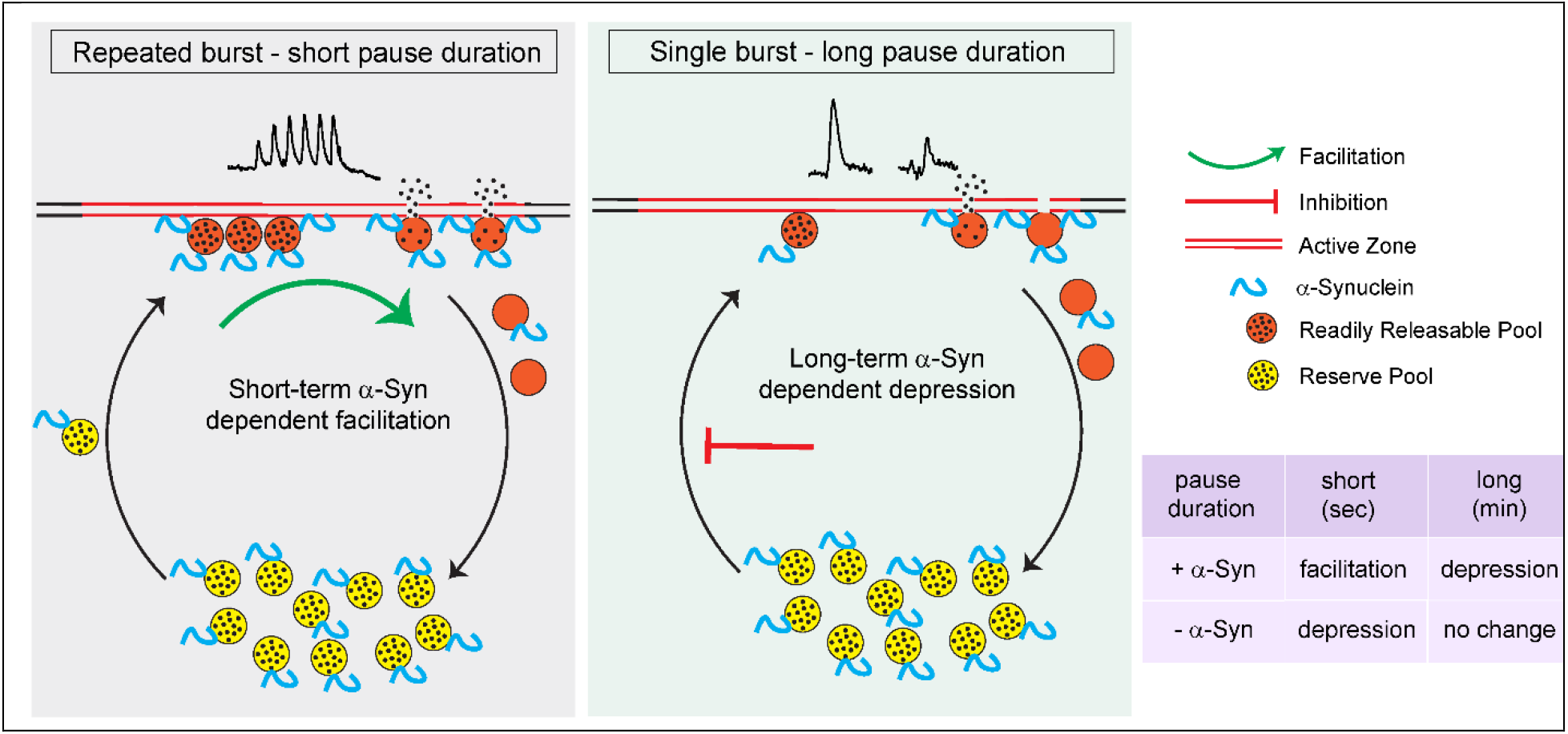
Model of α-Syn effects on dopamine release *in vivo*. In this model, the amount of release is rate-limited by the pool of synaptic vesicle/active zone complexes competent for fusion upon stimulation-dependent increase in calcium (the readily releasable pool (RRP), red). During the short intervals between repeated bursts (seconds; left), α-Syn enhances the dilation and collapse of synaptic vesicles, which in turn enhances the recovery of the active zones. Faster replenishment of the RRP allows larger number of synaptic vesicles to fuse in response to subsequent stimuli. In the absence of synuclein, active zones remain occupied for longer durations and the replenishment of the RRP is slower, resulting in presynaptic depression. With substantially longer intervals between bursts (minutes; right), the difference due to the rapid effects on synaptic vesicle dilation and collapse are negligible, revealing an inhibitory effect of α-Syn on a slower rate of transfer of vesicles from a reserve pool (yellow) to the RRP, leading to stimulation-dependent synaptic depression that is not observed in the absence of α-Syn.

While the α-Syn-dependent facilitation of DA release is consistent with an increase of competent synaptic vesicle/active zone complexes, there are alternative explanations, including effects on SNARE proteins that enhance docking or fusion (Burré et al., 2014; Hawk et al., 2019; Lou et al., 2017b) or enhancement of synaptic vesicle endocytosis (Vargas et al., 2014) and synaptic vesicle recycling. There is evidence from amperometric recordings that most dopamine synaptic vesicle release events may occur via a partial release of synaptic vesicle content (Staal et al., 2004) and if so, longer pore open times in the absence of synuclein expression could also enhance dopamine release. Unfortunately, quantal dopamine release events are too small to be measured *in vivo* with available technology.

We note that α-Syn- and activity- dependent presynaptic facilitation of DA release was not reported in a prior *in vivo* study (Yavich et al., 2004). This may be due to differences in the electrochemical detection technique (amperometry), site of electrical stimulation (medial forebrain bundle), stimulus paradigm (longer stimulation with less recovery time) or method of anesthesia (chloral hydrate, which can lead to silencing of both tonic and phasic activity by neurons (Koulchitsky et al., 2012)).

Consistent with prior reports (Abeliovich et al., 2000; Yavich et al., 2004) but not (Senior et al., 2008), we found that dopamine release evoked by single bursts was similar in the dorsal striatum of WT, SynTKO and α-SynKO mice, but decreased only in WT when stimulated by successive bursts separated by minutes-long pauses. The finding that synucleins can depress vesicle fusion in a calcium-independent manner appears consistent with our previous studies of the effects of synuclein overexpression on quantal catecholamine release in chromaffin cells (Larsen et al., 2006). This slow depression in WT became apparent at >20 minutes, with the 6^th^ burst releasing ~50% of initial levels. Surprisingly, this slow depression was not enhanced by increasing the number of stimuli during this period and does not appear to be due to tissue damage, as it was absent in the KO lines (Fig 3C and E).

The slow depression could be due to phosphorylated, aggregated or multimeric synucleins that form following activity-dependent dissociation from synaptic vesicle membrane (Fortin et al., 2005) and inhibit the mobility of synaptic vesicles that arrive over the course of minutes perhaps from “silent synapses” (Pereira et al., 2016), as the release of dopamine from *en passant* sites on striatal dopaminergic axons appears to be limited by the presence of active zone scaffolding proteins (Liu et al., 2018b). Alternatively, these forms may block SNARE function or other steps required for synaptic vesicle fusion (Hawk et al., 2019; Lou et al., 2017a).

### Summary of effects of synuclein expression on dopamine release in vivo

We find that the tonic firing of substantia nigra dopamine neurons is retained during the light anesthesia and that the levels of extracellular DA during this activity are beneath detection limits (<50 nM). As tonic firing is far slower than presynaptic recovery within bursts (~250 msec vs. < 10 msec), synuclein expression is expected to exert little effect on synaptic vesicle/active zone complexes during tonic dopamine release.

When a single burst exceeds ~ 5 pulses at 50 Hz, dopamine reuptake is saturated, and extracellular DA is linearly dependent on the number of pulses. However, the bursting eventually engages a slow depression of subsequent dopamine release, suggesting a decreased number of synaptic vesicle/active zone complexes.

During closely spaced repetitive bursts, there is an α-Syn-dependent facilitation of evoked dopamine release, consistent with an enhancement of competent synaptic vesicle/active zone complexes. The rapid facilitation can occur in tandem with the ongoing slow depression.

In summary, under conditions that emulate physiologically relevant bursting activity *in vivo*, α-Syn facilitates dopamine release. The effects of synucleins on dopamine transmission appear to be exquisitely tuned to inhibit rapid depression during repeated bursts, and could contribute to short-term plasticity that may underlie reinforcement and motor learning and action selection. Further research may indicate if this form of regulation occurs in other synapses that undergo prolonged bursting, such as those of the torpedo electric organ, neuromuscular junction, motor cortex, and the sympathetic nervous system.

## METHODS

### Mice

All experimental procedures followed NIH guidelines and were approved by Columbia University’s Institutional Animal Care and Use Committee. Data from male and female mice were combined since there were no differences in the experimental outcome between animal sexes. C57BL/6J (Stock No:000664), background controls (B6129SF2/J, Stock No: 101045) and α-Synuclein knockout (aSynKO, B6;129X1-Sncatm1Rosl/J, Stock No:003692) were acquired from Jackson Laboratories (Bar Harbor, ME, USA). The wild type (WT) mice consists of a pool of C57BL6/J and B6129SF2/J mice). Synuclein triple knockout (SynTKO) mice were obtained from the laboratory of Robert Edwards (UCSF, USA). All experiments were performed on 5-8-month-old mice.

### Surgery

Mice were anesthetized with isoflurane (SomnoSuite Small Animal Anesthesia System, Kent Scientific; induction 2.5%, maintenance 0.8–1.4% in O2, 0.35 l/min). A mouse was placed on a circulating warm water heating pad and head-fixed on a stereotaxic frame (Kopf Instruments, Tujunga, CA). Puralube vet ointment was applied on the eye to prevent cornea from drying out. Craniotomy (unilateral, right) was performed using a drill (0.8 mm) to target the specific region of interest with the following coordinates from mouse brain atlas (Franklin and Paxinos, 2008) – mm from Bregma - midbrain: AP = −2.9, ML=+1.0, DV=+4; Dorsal Striatum: AP=+1.2, ML=+1.3, DV=+3.1. Breathing and the depth of anesthesia (toe pinch) were constantly monitored.

### *In vivo* electrophysiology

Extracellular single-unit recordings of midbrain substantia nigra (SN) DA neurons were recorded from WT mice under isoflurane anesthesia (1-1.2%). We followed a previously described procedure (Subramaniam et al., 2014), where anesthetized mouse was head fixed on a stereotactic frame with continuous flow of isoflurane, followed by craniotomy to target midbrain as above (see Surgery). Glass electrodes (12–22 MΩ; Harvard Apparatus) filled with 0.5 m NaCl, 10mm HEPES, 1.5% neurobiotin (Vector Laboratories) were used for recording. An Ag/AgCl reference electrode was placed under the skin. Micromanipulator (SM-7, Luigs and Neumann) was used to lower the glass electrodes to the recording site. The spontaneous extracellular single-unit activity of SN DA neurons was recorded for 10 min using ITC-18 A/D converter (Heka, WinWCP software; sampling rate 11 kHz). The extracellular signals were amplified 1000× (ELC-03M, NPI Electronics), bandpass-filtered 0.3–5 kHz (single-pole, 6 dB/octave, DPA-2FS, NPI Electronics). Midbrain SN DA neurons were identified by the well-established electrophysiological characteristics - broad biphasic action potential (>1.2 ms), slow firing frequency (1–8 Hz) and their firing pattern (regular, irregular, and bursting; Grace and Bunney, 1984; Ungless et al., 2004). For the in vivo data analysis, the mean discharge frequencies were calculated and interspike interval histogram (ISIH,10ms bins) were plotted for every exported 600s spike-train. Coefficient of Variation (CV) is the measure of regularity of spiking, which was obtained by the ratio of standard deviation (SD) to the mean. The pattern of the in vivo spike-train was visually classified using the output from autocorrelation (measures ISIs to the nth order) histogram (ACH, 1 ms bins, smoothed with Gaussian filter (20ms)) (Perkel et al., 1967; Subramaniam et al., 2014; Tepper et al., 1995; Wilson et al., 1977) using R statistical computing (www.r.project.org).The observed in vivo firing patterns are: regular (pacemaker), irregular and bursty. The burst within the spike-train was identified according to the 80/160ms criterion (Grace and Bunney, 1984). A spike-train was classified as a regular if the ACH has a minimum of 3 consecutive oscillations with decreasing amplitude (the decrease in the amplitude is due to the finite timestamps in the spike-train i.e 600s). Irregular spike-train has an ACH with a plateau (equal probability of occurrence of specific ISI throughout the spike-train). Bursty pattern has an ACH showing a narrow initial peak (bursts) with a shallow trough followed by a broader peak (pause). Irregular bursty pattern has an ACH that shows an initial narrow and steep peak followed by a plateau. IgorPro 6.02 (WaveMetrics) software was used for the spike-train analysis.

### Juxtacellular labeling of single neurons *in vivo*

To map the anatomical localization of DA neurons, following extracellular single-unit recordings, the DA neurons were labeled with neurobiotin using juxtacellular *in vivo* labeling technique (Pinault, 1996). Microiontophoretic current was applied (1–10 nA positive current, 200ms on/off pulse, ELC-03M; EPC-10, NPI Electronics, as external trigger) via the recording electrode with continuous monitoring of the single-unit activity. The labeling event was considered successful when the firing pattern of the neuron was modulated during current injection (i.e., increased activity during on-pulse and absence of activity in the off-pulse) and the process was stable for minimum 25s followed by the spontaneous activity of the neuron after modulation. This procedure resulted in the exact mapping of the recorded DA neuron within the SN subnuclei (Franklin and Paxinos, 2008) along with its neurochemical identification using TH immunostainings.

### *In vivo* FSCV

Mice were anesthetized as described above and craniotomy was performed to target right midbrain and the striatum (see Surgery). A 22G bipolar stimulating electrode (P1 Technologies, VA, USA) was lowered to reach ventral midbrain (between 4-4.5mm). The exact depth was adjusted for maximal DA release. An Ag/AgCl reference electrode was placed under the skin via a saline bridge. A carbon fiber electrode (5 μm diameter, cut to ∼150 μm length, Hexcel Corporation, CT, USA) was used to measure the evoked DA release from the dorsal striatum. To verify electrode positions in FSCV experiments, the carbon fiber electrode was briefly dipped into DiI solution (1:1000, ThermoFisher, USA) and air dried. This electrode was lowered to the described coordinates. The stimulating electrode position was identified using the electrode track in the brain tissue. Constant current (400μA) was delivered using an Iso-Flex stimulus isolator triggered by a Master-9 pulse generator (AMPI, Jerusalem, Israel). A single burst stimulation consisted of 30 pulses at 50Hz (0.6s). Repeated-burst stimulation consisted of six 30 pulse/50 Hz bursts delivered at 5 s interval. Electrodes were calibrated using known concentration of DA in ACSF. Custom-written procedure in IGOR Pro was used for the data acquisition and analysis.

### *In vivo* fiber photometry

Mice were anesthetized using isoflurane and prepared for surgery and craniotomy targeted to the midbrain (see Surgery). A Nanoject was used to inject 100nl of AAV9-Flex-GCamp6f (vendor) at a rate of 18.4 nl/min, followed by 5 minutes wait period before removing the needle. Mice were left to recover for three weeks to allow for GCamp6f expression. On the day of the experiment mice were anesthetized and craniotomy was performed to target midbrain and the striatum (see Surgery). An optic fiber (Doric, 0.22NA, 300um diameter, 3.5mm length) was lowered in the dorsal striatum, while stimulating electrode was placed to the ventral midbrain. Doric photometry system (Doric Lenses Inc, Canada) was used for light excitation and detection. Two LED were used, one at wavelength 405nm for background fluorescence, and a second one at 465nm to excite GCamp6f. The same fiber was used to detect the light emission. Doric Neuroscience Studio software was used to record the data. Calcium traces were analyzed with custom-written codes using Python and IgorPro6.2.

### Histology and imaging

For immunocytochemical detection, mice were transcardially perfused with ice-cold paraformaldehyde (PFA, 4%) and the brains were isolated and stored in PFA overnight. Next, brains were sectioned into 50 μm (for TH and biotin staining) or 100 μm (for DiI visualization) slices using Leica VT2000 vibratome (Richmond, VA). Free-floating sections were rinsed with PBS (3×10min) followed by 1 hour blocking at room temperature using 10% NDS (normal Donkey Serum) in PBS/TX (0.1-0.3%). Sections were then incubated overnight with primary antibody against tyrosine hydroxylase diluted in 2% NDS in PBS/TX (1:1000 - rabbit anti-TH, Millipore AB152). The sections were rinsed with PBS (3×10min) and incubated for 6 hours with secondary donkey anti-rabbit 488 antibodies (1:1000, ThermoFisher Scientific, A11006) and streptavidin 594 (1:750, Invitrogen, S32356) diluted in 2% NDS in PBS/TX. The sections were washed with PBS (3×10min), arranged along the caudo-rostral axis, mounted onto Superfrost plus microscope slides (Fisher Scientific, USA) and allowed to air-dry. Once the sections were partially attached to the glass, the coverslips were mounted using an anti-fading mounting medium (Vectashield, Vector Laboratories, Burlingame, USA). The sections were visualized using confocal laser scanning microscopy (CLSM) (Leica TCS SP8, 5x, 10x and 60x objectives) operated by Leica LAS X Core software.

### Statistical Analysis

The data in the figures are represented as mean with standard error of mean (S.E.M). Grouped data (with one or two different conditions) were analyzed using repeated measures one/two-factor within-subject analysis of variance (1-way or 2-way ANOVA, mixed effects model), followed by Bonferroni’s multiple comparisons test. The dependent measures were the: burst numbers and genotypes. Significance was assumed for all values where p<0.05. The values for each analysis (mean+_SEM, F and degree of freedom (DF) values are presented in the text). GraphPad Prism Software (GraphPad Software, San Diego, CA) was used for the data plots and statistical analysis.

## Acknowldgements

We gratefully acknowledge Rodrigo Espana and his team (Drexel University, PA) for their help in setting up the *in vivo* FSCV technique. We thank Dr. Mark Wightman for valuable comments on the manuscript and insightful feedback and to Shashaank N. for proofreading. Supported by NIH R01DA04718 and the JPB Foundation.

## Author Contributions

M.S, E.V.M, D.S: conceived and designed the research; M.S, S.C: performed the experiments; M.S: analyzed data; R.H.E: provided SynTKO animals and contributed to the Discussion; M.S and D.L.S wrote the manuscript; M.S, S.C, R.H.E, E.V.M, D.S: reviewed and edited the manuscript.

---

**Figure S1:**
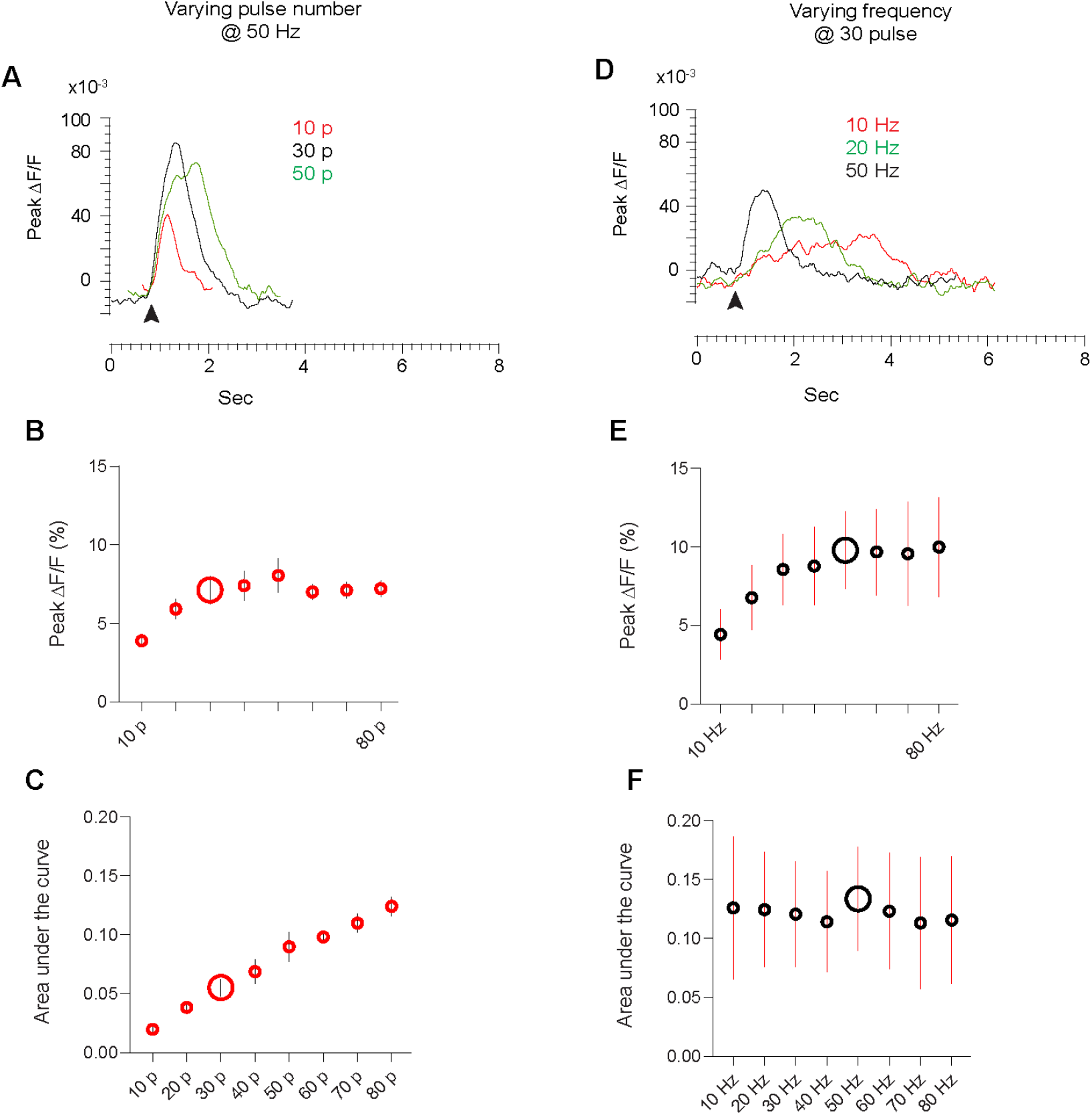
Analysis of in vivo GCaMP6f response to electrical stimulation. **A**, Evoked calcium response to electrical stimulation at constant frequency (50 Hz) and varying pulse number (10 pulse – red; 30 pulse – black; 50 pulse – green) under isoflurane anesthesia. Arrowhead indicates the initiation of electrical stimulation. **B**, Peak GCaMP6f signal (ΔF/F) at varying pulse number (10-80 p) at a constant stimulation frequency. Note the plateau in the signal peak height from 30 p. **C**, Area under the curve (AUC) measurements for the evoked calcium signals at different pulse number (10-80 p). Note the linear increase in the AUC (despite the lack of change in peak height). **D**, Evoked GCaMP6f response to electrical stimulation at constant pulse number (30 pulses) at a range of stimulus frequencies (10 Hz – red; 20 Hz – green; 50 Hz – black). Arrowhead indicate the initiation of electrical stimulation. **E,** Peak calcium signal (ΔF/F) at varying stimulation frequency (10-80 Hz) at a constant pulse number. Note the plateau in the GCaMP6f signal from 50 Hz. **F**, Area under the curve (AUC) measurements for the evoked calcium signals at different stimulation frequencies (10-80 Hz). Note that change in stimulation frequency has no effect on the AUC. Enlarged circles represent the chosen stimulation parameters. Error bars represent SEM (n= 4 mice).

